# The tree cricket ear is a highly phase sensitive biomechanical interferometer

**DOI:** 10.1101/2025.10.28.684507

**Authors:** Emine Celiker, Gregory Patrick Sutton, Natasha Mhatre

## Abstract

We show that tree crickets localize the sound by having an auditory system that first splits the soundwave into two parts, sending one wave to an Anterior tympanic membrane, and sending the second to a Posterior tympanic membrane. The membranes then carry the split waves to a common point, the Tracheal wall, where the re-assembly of the wave causes mechanical interference that is sensitive to microsecond time delays. This interference mechanically transduces the time delay into a simple lateral strain at the Tracheal wall. This principle, interferometry, is usually thought of in terms of light (Michaelson and Morley, 1887), and can measure extremely small time-delays. Tree crickets use the same principle as the light interferometer, except with sound, to measure extremely precise locations.

---

Finding a singing cricket in a living room is infuriatingly difficult. While large vertebrates, like us, can determine the direction of sound from several features that are altered as the sound hits the left and right ears (Grothe et al., 2010), resolving this precisely requires the nervous system to react to small (hundreds of microseconds) time differences between the ears (von Kriegstein et al., 2008). For crickets, using this mechanism is even more difficult because their small size would require their nervous system to differentiate between even impossibly smaller (microsecond) time differences; and yet female crickets have no trouble locating singing males (Mhatre and Balakrishnan 2008), even when one ear is removed (Huber et al., 1984; Kohne et al., 1992). We will show that tree crickets (*Oecanthus henryi*) accomplish this task by using an elegant method of mechano-acoustic interferometry (Figure 1D,E).

**Figure 1.**
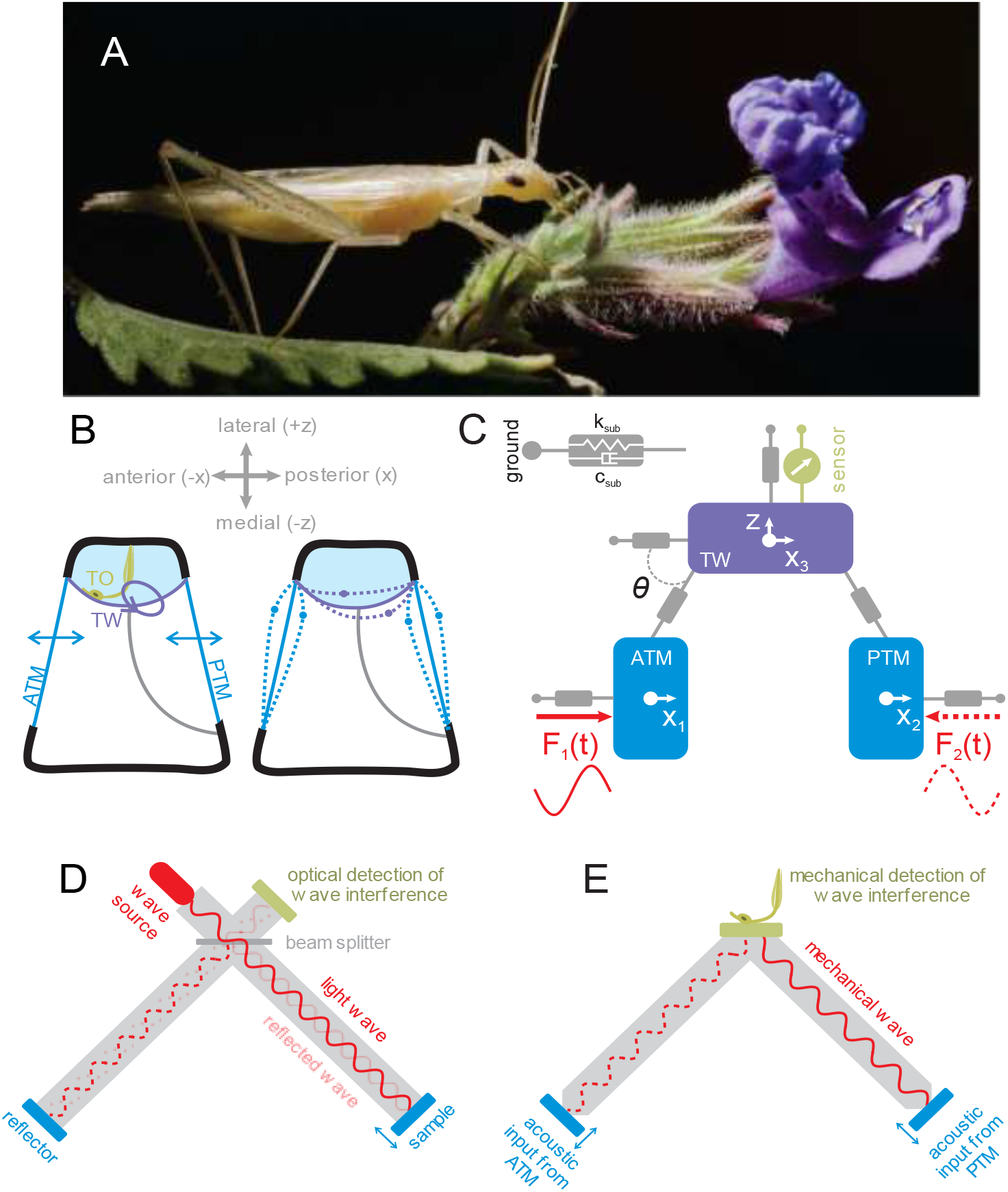
The tree cricket ear anatomy. A. The tree cricket *Oecanthus henryi*. B. A cross-sectional sketch of the tree cricket *Oecanthus henryi* ear near the tympanal end of the acoustic trachea. ATM = Anterior Tympanic Membrane, PTM = Posterior Tympanic Membrane, TO = Tympanal Organ, TW = Trachea Wall. C. The fully-coupled mechanical system representing the tracheal wall (TW) displacement. The tracheal wall (stiffness k_TW_=20.62 N/m, damping c_TW_=0.00047, and mass M_TW_=2.8×10^-10^ kg) is acted upon by the anterior tympanic membrane (k_ATM_=41.23 N/m, damping c_ATM_=0.00047, and mass M_ATM_=1.6×10^-10^ kg), and posterior tympanic membrane (k_PTM_=41.23 N/m, damping c_PTM_=0.00047, and mass M_PTM_=1.4×10^-10^ kg). The angle between the tympana and tracheal wall *θ* = 60° (Mhatre et al., 2009). F_1_(t) and F_2_(t) represent the harmonic forces of equal magnitude and a frequency of 3 kHz acting on the tympana. D. A schematic of the light interferometer mechanism. E. The schematic of the tree cricket ear acting as a biomechanical interferometer.

Tree crickets (in the suborder *ensifera*) (Figure 1A) possess ears located in the tibia of their forelegs (Michelsen et al., 1994; Mhatre et al., 2009). The ear in each leg has two tympana: an anterior tympanic membrane (ATM) and a posterior tympanic membrane (PTM). These tympana are backed by an air-filled tube called the acoustic trachea, with an opening, the acoustic spiracle, located on the thorax. Each tympana receives multiple inputs of the same sound: directly to the external side, and through the acoustic trachea (Michelsen et al., 1994; Michelsen, 1998). Both membranes attach ventrally to a common cuticular structure, the tracheal wall (TW), with sound waves being carried by both membranes to this common point (Figure 1B). The tracheal wall is also where the sensory neurons are located (Field and Matheson, 1998; Mhatre et al., 2021). While the geometry of the ear does lead to a direction dependent intra-aural time delay (Pulver et al., 2022), this time delay (on the order of microseconds) is orders of magnitude smaller than the time delays detected by vertebrates. Larger crickets are thought to solve this problem using a four-input pressure difference system (Larsen and Michelson, 1978), but for even smaller insects this requires novel solutions. Flies solve this by using an independently evolved inter-tympanal mechanical linker (Mason et al., 2001), but it is unclear how smaller crickets like tree crickets solve the problem of sound localization.

We have mechanically approximated the tree cricket ear with a three-element mechanical model: 1) an anterior tympanic membrane, which receives one sound input, 2) a posterior tympanic membrane, which receives a second sound input, 3) a tracheal wall, which is acted upon through the displacement of each membrane (Figure 1B). The specific mass, stiffness, and damping parameters used, and the mathematical representation of the lumped element model are detailed in supplementary materials. This three-element model mechanically recreates the acoustic response of *O. henryi* tracheal wall (Figure 1C).

The lumped element model shows that the key to this system is the individual membranes carrying the soundwaves to the common point, the tracheal wall, where the interference between the two soundwaves causes the strains at the tracheal wall to differ, depending on a phase difference between the sound waves hitting each membrane (Figure 2A,C and supplemental movie 1). A zero degree phase difference is transduced into vertical strains in the tracheal wall, a 180 degree phase difference into horizontal strains, and intermediate phase differences into elliptical strains (Figure 2A and supplemental movie 1). The acoustic inputs resulting in elliptical movements have been observed previously in both insects and mammals (Stephen and Bennet-Clarke, 1982; Vavakou et al., 2024; Frost et al. 2025) with Bennet-Clark commenting “it would appear at first sight that the two drives will interact to reduce the sharpness of tuning of each other”; we show here that these ellipses are tied to the phase differences in the sound (Figure 2A,C), and thus can be used to precisely determine the direction of the sound source. It is this phase-dependent interference between the two sound waves detected by the nervous system – making the tree cricket ear a biological example of a mechano-acoustic phase interferometer akin to that developed for light in Michelson and Morley (1887) (Figure 1 D,E).

**Figure 2.**
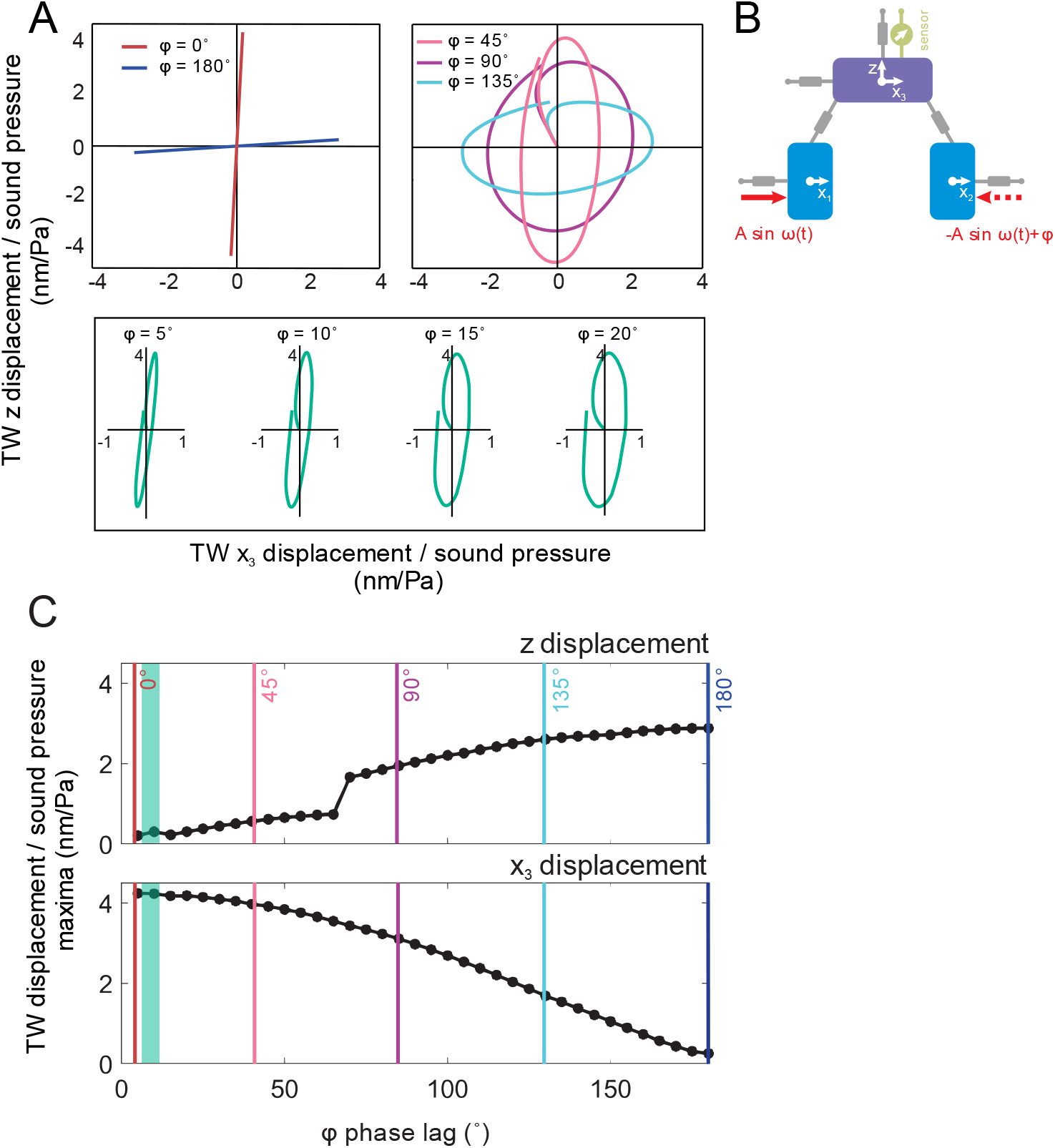
The displacement sensitivity of the tracheal wall at the calling song frequency 3 kHz (at 24 °*C*) of *Oecanthus henryi*. Phase lag refers to the phase difference between the forces of same magnitude activating the anterior tympanic membrane (ATM), and the posterior tympanic membrane (PTM). A. The displacement pattern of the tracheal wall when the ATM and PTM are acted on by forces *in-phase* (phase lag=0°) and *out-of-phase* (phase lag=180°) (top left); The elliptic motion of the tracheal wall for large phase differences between the ATM and PTM (top right); The elliptic motion of the trachea wall for small phase differences between the ATM and PTM (bottom). B. The lumped element model representing the trachea wall displacement. C. The displacement magnitude of the tracheal wall in the x_3_- and z-directions with respect to the phase lag of the forces acting on the ATM and PTM.

This biomechanical phase interferometer requires a small number of simple components: 1) two similarly-sized membranes transducing disparate vibrational inputs to a common point moving at similar levels, 2) a system that allows for sound direction to create a phase difference between the soundwaves hitting each membrane, and 3) mechanosensory neurons at that common point (the tracheal wall) that respond preferentially to a single direction of movement.

The first of these components is a feature common to many Ensiferan species and its functional significance was hitherto unappreciated. For instance, bush crickets (fellow *ensiferans*), have a very similar ear morphology consisting of two membranes carrying the sound to a common point (Montealegre-Z and Robert, 2015). For component two, small phase differences between closely placed tympana may be created due to diffraction by the body of the animal, ear anatomy, and the substrate it stands on. Indeed, a directionally dependent phase difference in the sound hitting the ATM and PTM has been observed in bush crickets (Pulver et al., 2022). Such a difference is accentuated by the geometry of the tracheal branches carrying the sound to each membrane (Veitch et al., 2021). Moreover, similar elliptical movements in the tracheal wall have been observed in tree crickets (Mhatre et al, 2021) and also in bush crickets (Vavakou et al., 2024), strongly suggesting that this phase interferometry method is widespread among Ensiferans. Finally, it is also possible that mechanosensory neurons respond preferentially to certain axes of stimulation (Yarger and Fox, 2018; Chaiyasitdhi et al., 2025). The simple components required for such a system also lead to the possibility that many hearing systems in small animals may be similar.

This simplicity provides a huge number of parameters that can then be further tuned to specialize different Ensiferan ears for different inputs. Membrane geometry (Mhatre et al., 2009; Rajarman et al., 2013), membrane mechanical properties (Vincent and Wegst, 2004), membrane anisotropy (Mhatre et al., 2009; Mhatre et al., 2012; Malkin et al., 2014), acoustic trachea geometry (Schmidt and Römer, 2013; Veitch et al., 2021; Celiker et al., 2022), fluid properties within the auditory vesicle (Veitch et al., 2021; Lomas et al., 2012), or mechanisms for active amplification (Mhatre and Robert, 2013) could all be modified to specialize the interferometer for specific sound inputs.

Moreover, as this system is sensitive to the phase of the sound (as opposed to the time delay) - the higher the frequency of the sound, the easier it is for the system to mechanically transduce the precise direction of the sound’s source. This would have obvious utility in detecting bats (which have call frequencies in excess of 60kHz, Pulver et al., 2022) as well as detecting the song in ultrasonic crickets (Robillard et al., 2013).

We have trouble locating that singing cricket because of its tonal nature, and our nervous system has difficulty resolving the minute time delays in such signals. Tree crickets, on the other hand, pinpoint the sound using mechano-acoustic phase interferometry, letting the tympanal membranes and the tracheal wall precisely tell them where a singing conspecific is.

## Acknowledgements

NM was partially supported by the Canada Foundation for Innovation (CFI4748) and the Ontario Research Fund, NSERC Discovery (Grant no. 687216), early career supplement (675248) and an NSERC Canada research chair (Grant no. 693206) and the Western SEED grant. GPS was supported by a University Research Fellowship from the Royal Society, UK (UF130507)

