## Supplementary Materials for "The tree cricket ear is a highly phase sensitive biomechanical interferometer"

#### S1 Mechanism of tree-cricket tympanum

To obtain a qualitative understanding of the mechanical properties of the tree cricket *O. henryi* tympana, we modelled the ATM using a lumped element approach (Figure S1). The model is characterised by an effective mass ($M$), effective stiffness ($k$), and effective damping ($c$).

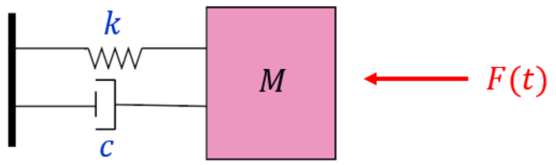


Figure S1: The mass-spring-damper system representing the *Oecanthus henryi* tympanum mechanism. $M, k$ and $c$ are the mass, stiffness and damping parameters of the tympanum, respectively.

The tympanal displacement magnitude was obtained from the solution to the equation of motion in the frequency domain,

$$(-M\omega^2 + jcw + k)X(\omega) = F(\omega), \tag{1}$$

which is the mathematical representation of the system given in Figure S1. The dependent variable $X(\omega)$ is the complex displacement with respect to the angular frequency ($w = 2\pi f, f$ =frequency), $j = \sqrt{-1}$, and the externally applied force $F(\omega) = F_0$ represents the sound stimulus. The magnitude of the applied mechanical force was taken as $F_0 = P_0 \times A$, where $P_0 = 0.04$ Pa and $A$ is the cross-sectional area of the membrane (Table S1), which is equivalent to an acoustic pressure of magnitude $P_0 = 0.04$ Pa acting on the tympanic membrane [1]. The solution was considered in the frequency range 2-7 kHz.

The analytical solution to equation (1) takes the well-known form ([2], Ch. 1, pp.10)

$$X(\omega) = \frac{P_0 \times A}{j\omega(c + j(\omega M - k/\omega)}, \tag{2}$$

which gives the displacement magnitude as

$$|X(\omega)| = \frac{P_0 \times A}{\omega\sqrt{c^2 + (\omega M - k/\omega)^2}}, \tag{3}$$

with the corresponding phase angle

$$\Theta = \tan^{-1}\left(\frac{c}{\omega M - k/\omega}\right). \tag{4}$$

Next, the parameter values of $M$, $k$ and $c$ that fit the known *O. henryi* ATM properties were considered. The total mass of the tympanum was calculated using the formula

$$M_{tot} = A \times h \times \rho, \tag{5}$$

where $A$ = cross-sectional area, $h$ = thickness and $\rho$ = density of ATM, whose values are listed in Table S1. The ATM was assumed to have an elliptic cross-sectional area, and its dimensions were taken as measured in [3] and [4] for *O. henryi*. The density was taken as the mean density value of insect cuticle due to the narrow variation between measurements (cf. [5]). However, the LDV measurements of the *O. henryi* ATM demonstrated that only small portions of the membrane are set in motion to sound, with maximal deflection occurring in the central region of the membrane (the fundamental mode) [3]. Making the simplifying assumption that the active region is a uniform clamped circular membrane vibrating at the fundamental mode, we accordingly adopted an effective mass as a fraction of its total mass, namely $M = 0.27M_{tot}$ [6].

| Parameter | ATM |
|---|---|
| Length | $9 \times 10^{-4}$ m |
| Width | $3.5 \times 10^{-4}$ m |
| Cross-sectional area ($A$) | $2.5 \times 10^{-7}$ m$^2$ |
| Thickness ($h$) | $2 \times 10^{-6}$ m |
| Density ($\rho$) | 1200 kg/m$^3$ |
| Total Mass ($M_{tot}$) | $5.9 \times 10^{-10}$ kg |
| Effective Mass ($M$) | $1.6 \times 10^{-10}$ kg |

Table S1: ATM dimensions of tree cricket *Oecanthus henryi* used to obtain the effective mass, $M$.

For the value of the stiffness parameter $k$ of the ATM, we made the simplifying assumption that the tympanum is isoptropic and homogeneous. As the ATM response to sound is flat in the frequency range 2-7 kHz [3], [7], it is clearly stiffness dominated in this frequency range ([2], Ch.1, pp.18). Utilising this static force-to-displacement relationship, we obtain the effective stiffness from the analytical solution (3). For a stiffness dominated system, the dominant term in the denominator's square root of formula (3) becomes $k/\omega$. As seen from the LDV recordings presented in [3] and [7], in the frequency range 2-7 kHz the sensitivity of the ATM is ∼6 nm/Pa. Using this, the effective stiffness $k$ is obtained from the relation

$$6 \times 10^{-9} = \frac{|X(\omega)|}{P_0} \sim \frac{A}{k}. \tag{6}$$

Rearranging, we obtain the effective stiffness as

$$k \sim \frac{2.474 \times 10^{-7}}{6 \times 10^{-9}} = 41.23 \text{ N/m}.$$

A validation of the stiffness value is given in Section S1.1.1.

Finally, as the tree cricket tympanum is expected to be viscoelastic [8], we obtained a frequency dependent effective damping parameter using parameter sweeps based on equation (3) and a comparison with the LDV data [7] (see Section S1.2.2).

The analytical solution described above was used to calculate the passive tympanal response to sound pressure in the frequency domain, within the range 2-7 kHz. Figure S2 shows a comparison between numerical results and the LDV recordings [7]. Consistent with experimental recordings, the analytical solution demonstrated a flat response of the ATM to the sound stimulus across the studied frequency range. The relative percentage error of ATM sensitivity between the mean experimental and numerical data was 11.5% (Section S1.1.2). For a direct comparison between the experimental and numerical phase data, the numerical data was calibrated by a magnitude of 20° to account for the fixed time delay in the experimental setup that is not available in the analytical solution. For the frequency range 2.5-7 kHz, the numerical phase calculations remained within the standard deviation bounds of the experimental phase recordings, and showed a maximum absolute difference of 28° with the mean LDV recordings (Section S1.1.2). In addition, both the experimental and numerical phase data demonstrated a similar, negative slope. Hence, the analytical solution for the ATM sensitivity and phase gave a good match with the LDV recordings.

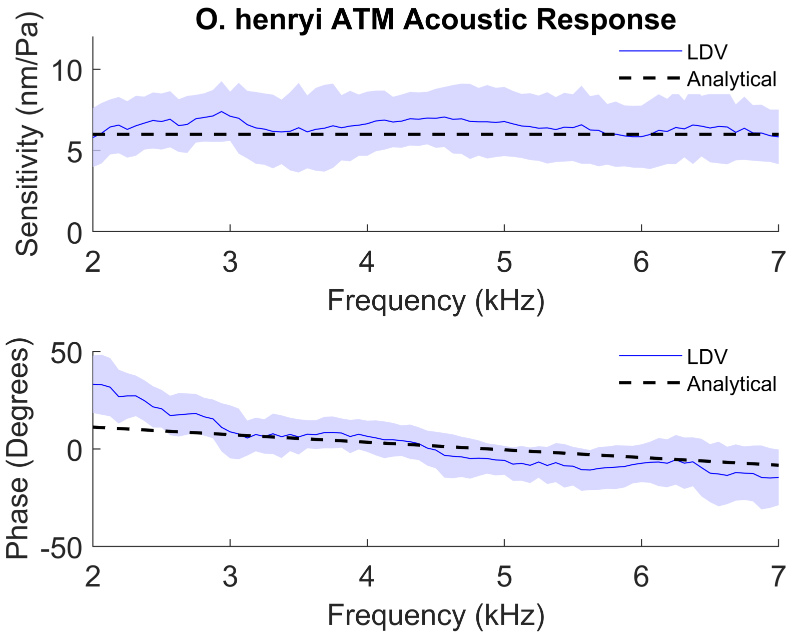


Figure S2: Comparison of the analytical solution and laser doppler vibrometer (LDV) recordings of tree cricket tympanal displacement magnitude (top figure) and phase (bottom figure) from [7]. Navy line shows the mean and shaded zone indicates ±1 SD of the LDV data from $n = 12$ animals.

#### S1.1 Tympana Material Parameters

##### S1.1.1 Tympanum effective stiffness validation

As a validation for the effective stiffness value calculated for the anterior tympanic membrane (ATM), we considered the static stiffness of an idealised tympanum - a clamped circular plate. The static stiffness was calculated using the formula

$$k_s = \frac{F}{\delta},$$

where $F$ =is the total force applied to the plate , and

$$\delta = \frac{Pr^4}{64D}$$

is the maximum deflection of the plate, with $P$=pressure per unit area, $r$ =plate radius and $D$ =bending stiffness ([9], Table 11.2, Condition 10b). The bending stiffness is calculated as $D = \frac{Eh^3}{12(1-\nu^2)}$, where $E, h$ and $\nu$ are the Elastic modulus, thickness and Poisson's ratio, respectively, as listed in Table S2. The thickness is the same as the tree cricket ATM thickness [3]. Since the tree-cricket tympana are elliptic, the plate radius was taken as the averaged major and minor axes of the measured dimensions [3]. The radius was then multiplied by 0.27 to reflect that only a small part of the plate is active [6]. Poisson's ratio was taken as a value generally used for insect tympana [10]. The Elastic modulus of tree cricket tympanum is not known, hence we carried out parameter sweeps of the static plate stiffness within known bounds of insect cuticle, [5], until there was a good match with the effective stiffness used for the lumped element model.

Table S2: Parameter values used to calculate the static stiffness of an idealised tympanum.

| Parameter | Value |
|---|---|
| Thickness ($h$) | 2 $\mu$m |
| Radius ($r$) | 84.375 $\mu$m |
| Poisson's Ratio ($\nu$) | 0.3 |
| Elastic Modulus ($E$) | 2 GPa |

Substituting the values given in Table S2 into the formula, and noting that total force $F = P\pi r^2$ [1], we obtained $k_S = 41.38$ N/m. Hence, the stiffness of the idealised tympanum compared remarkably well with the effective stiffness of the tree cricket ATM ($k = 41.23$ N/m).

##### S1.1.2 Tympanum damping

To calculate the frequency dependent damping of the tree cricket ATM, the analytical solution of the equation of motion was utilised. For this, we considered equation (3) which represents the displacement magnitude as obtained from the analytical solution. Rearranging this equation, we obtain the damping as

$$c(\omega) = \sqrt{\frac{A}{\omega}\frac{P_0}{|X(\omega)|} - (\omega M - k/\omega)^2}. \quad (7)$$

For consistency with the effective stiffness calculation, a tympanal sensitivity of $\frac{|X(\omega)|}{P_0} = 6$ nm/Pa was adopted from the experimental LDV recordings [7]. Substituting this into equation (7) gives

$$c(\omega) = \sqrt{\frac{A}{(6 \times 10^{-9})\omega} - (\omega M - k/\omega)^2}. \quad (8)$$

Equation (8) was used as a starting point for the effective damping parameter sweeps. For this, the constant multiples of equation (8) were applied to calculate the ATM displacement magnitude and phase while keeping the effective mass and stiffness fixed. The obtained results were first compared with the LDV data [7] schematically.

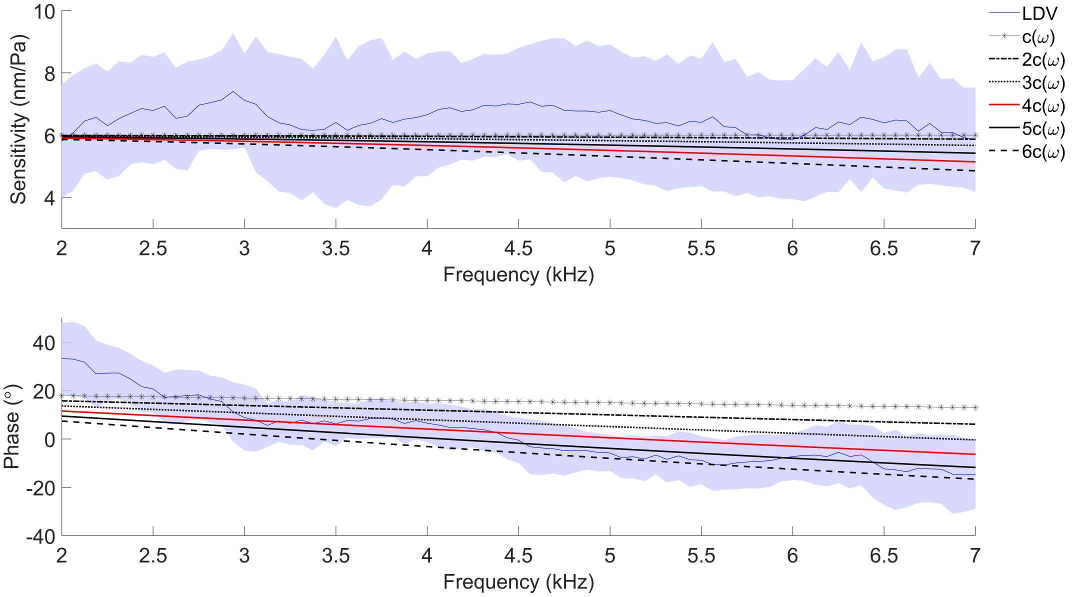


Figure S3: A comparison of the numerical and experimental ATM displacement magnitude (top figure) and phase (bottom figure), for constant multiples of the damping parameter $c(\omega)$.

As demonstrated from Figure S3, the numerical sensitivity calculations remained within the standard deviation bounds of the LDV recordings. To obtain the best fit, the percentage errors were calculated using the relative error formula

$$RE = \frac{1}{N}\sum_{i=1}^{N}\frac{|y_i - \hat{y}_i|}{|y_i|}100,$$

where $y_i, \hat{y}_i$, $i = 1, 2, \cdots N$ are the mean experimental and numerical data, respectively, and $N$ is the number of data points. The obtained results are presented in Table S3.

Table S3: Relative percentage error (RE) between the mean LDV recordings [7] and numerical results for sensitivity comparison.

| | RE (%) |
|---|---|
| $c(\omega)$ | 7.7 |
| $2c(\omega)$ | 8.4 |
| $3c(\omega)$ | 9.7 |
| $4c(\omega)$ | 11.5 |
| $5c(\omega)$ | 13.5 |

For a comparison between the experimental and numerical phase data, despite a calibration of 20° of the numerical data to account for differences with the experimental set-up, the results fell out of the standard deviation bounds for all considered damping parameters (Figure S3). Nevertheless, for $4c(\omega)$, the numerical phase data was within bounds for 2.5 - 7 kHz for the 2- 7 kHz frequency range considered. Since the numerical phase data had regions out of the standard deviation bounds, rather than calculating the percentage error, we calculated the maximum absolute difference between the two data sets. The obtained results did not show a significant difference between the considered damping parameter values (Table S4).

Table S4: Maximum absolute difference between the phase data of the mean LDV recordings ($E_p$) [7], and the numerical results ($N_p$).

| | $\max \lvert E_p - N_p \rvert$ (°) |
|---|---|
| $c(\omega)$ | 27.8 |
| $2c(\omega)$ | 24.2 |
| $3c(\omega)$ | 26.1 |
| $4c(\omega)$ | 28.0 |
| $5c(\omega)$ | 29.7 |

Based on the schematic investigation (Figure S3) and the following percentage error calculation for sensitivity (Table S3) and maximum absolute difference for the phase data (Table S4), the frequency dependent parameter values were adopted as $4c(\omega)$. The range of values of $4c(\omega)$ is given in Figure S4.

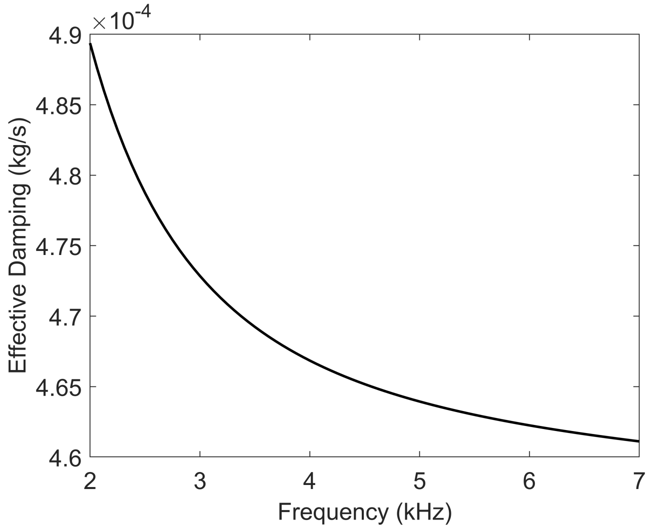


Figure S4: The frequency dependent effective damping values.

### S2 A fully-coupled lumped element model of trachea wall displacement

The OCT recordings of the tree-cricket ear by Mhatre et al. [11] showed that the displacement of the trachea wall was primarily guided by the tympanic membranes through their mechanical coupling. Based on this, we developed a lumped-element model of the tracheal wall displacement during passive state through its coupling to the ATM and PTM (Main text, Figure 1c). The displacement was considered in the time domain. The parameters $M_{-}, c_{-}$ and $k_{-}$ denoted on the model are the effective mass, stiffness and damping parameters, respectively, whose subscripts ATM, PTM and TW show which part of the tree cricket ear they belong to. The angle $\theta$ is for the natural slant of the tympana, and has a magnitude of $\theta = 60°$ [3]. The system is excited by the sound stimuli

$$F_1(t) = F_0 \cos(\omega t) \text{ and } F_2(t) = F_0 \cos(\omega t + \varphi)$$

acting on ATM and PTM, respectively, where $F_0 = 40$ mPa is the sound stimulus amplitude, $\omega = 2\pi f$ is the angular frequency with $f$ =3 kHz, and $\varphi$ is the phase angle.

The mathematical equivalent of the system is given in Supplementary Materials, Section S2.1.

The ATM parameter values were taken the same as in Section S1. Similar to the ATM, the PTM was also assumed to have an elliptic shape, with its precise maximum width and length of $3.5 \times 10^{-4}$ and $8 \times 10^{-4}$, respectively [3]. The mass of PTM was calculated as for ATM (see Section S1). Since the LDV recordings of the PTM sound response given by Mhatre et. al [3] demonstrated a flat response with a sensitivity equal to ATM, we took the effective stiffness $k_{PTM}$ to be the same as $k_{ATM}$. Likewise, $c_{ATM}$ and $c_{PTM}$ were obtained as in Section S1 at the considered frequency $f = 3$ kHz. The precise parameter values used for the ATM and PTM are listed in Table S5.

Although the empirical measures of the *O. henryi* material parameters are not available, detailed anatomical measurements of the related field cricket *Gryllus bimaculatus* showed a comparable thickness between the PTM and the TW [13]. Based on this assumption, we took the *O. henryi* TW thickness to be the same as the thickness of its PTM, namely 2$\mu$m. The length of the *O. henryi* TW was measured as 130 $\mu$m ([3], Figure 1(g)). As we are interested in the region of TW connected to the tympana, the width was taken as the width of the ATM, namely 900 $\mu$m. Hence, assuming a rectangular cross-sectional area for TW and using formula (5), the effective mass of the TW was calculated as

$$M_{TW} = 2.808 \times 10^{-10} \text{ kg}.$$

The effective damping parameter was taken to be the same as for ATM and PTM in both $x-$ and $z-$directions. For the remaining unknown parameter, the effective stiffness of the TW, we used parameter sweeps by comparing the numerical TW sensitivity with the OCT recordings of the tree cricket species *Oecanthus californicus*, whose passive state tympana also gave a flat response to sound within the measured frequency range 3-4.5 kHz, with a comparable sensitivity to *O. henry* [11] (see Section S2.2). The parameter values adopted for TW are listed in Table S5.

Table S5: Anterior tympanic membrane (ATM), posterior tympanic membrane (PTM) and trachea wall (TW) material properties used in the mathematical model.

| Parameter | ATM | PTM | TW |
|---|---|---|---|
| Mass (kg) | $1.6 \times 10^{-10}$ | $1.4 \times 10^{-10}$ | $2.8 \times 10^{-10}$ |
| Stiffness (N/m) | 41.23 | 41.23 | 20.62 |
| Damping (kg/s) | 0.00047 | 0.00047 | 0.00047 |

Although recorded using a different tree cricket species (*Oecanthus californicus*), a comparison between the OCT measurements of TW displacement magnitude given in [11] for the passive state, and the numerical results (Main text, Figure 2) showed a notable match, validating the accuracy of the lumped element model.

#### S2.1 Mathematical representation of tree cricket ear biomechanism

The mechanical system in Figure 2b (Main text, Section 2.2) is mathematically described by the following system of second-order ordinary differential equations.

$$
\begin{aligned}
M_{ATM}x_1''(t) = F_1(t) \quad &+ \quad k_{ATM}\left(\sqrt{(x_3(t)-x_1(t))^2+z(t)^2}-\sqrt{(x_3(0)-x_1(0))^2+(z(0))^2}\right)\cos(\theta) \\
&+ \quad c_{ATM}\frac{d}{dt}\left(\sqrt{(x_3(t)-x_1(t))^2+z(t)^2}\right)\cos(\theta)+k_{ATM}(x_1(0)-x_1(t))-c_{ATM}x_1'(t) \\
M_{PTM}x_2''(t) = -F_2(t) \quad &- \quad k_{PTM}\left(\sqrt{(x_2(t)-x_3(t))^2+z(t)^2}-\sqrt{(x_2(0)-x_3(0))^2+(z(0))^2}\right)\cos(\theta) \\
&- \quad c_{PTM}\frac{d}{dt}\left(\sqrt{(x_2(t)-x_3(t))^2+z(t)^2}\right)\cos(\theta)+k_{PTM}(x_2(0)-x_2(t))-c_{PTM}x_2'(t) \\
M_{TW}x_3''(t) = \quad &- \quad k_{ATM}\left(\sqrt{(x_3(t)-x_1(t))^2+z(t)^2}-\sqrt{(x_3(0)-x_1(0))^2+(z(0))^2}\right)\cos(\theta) \\
&- \quad c_{ATM}\frac{d}{dt}\left(\sqrt{(x_3(t)-x_1(t))^2+z(t)^2}\right)\cos(\theta) \\
&+ \quad k_{PTM}\left(\sqrt{(x_2(t)-x_3(t))^2+z(t)^2}-\sqrt{(x_2(0)-x_3(0))^2+(z(0))^2}\right)\cos(\theta) \\
&+ \quad c_{PTM}\frac{d}{dt}\left(\sqrt{(x_2(t)-x_3(t))^2+z(t)^2}\right)\cos(\theta)-k_{TW}x_3(t)-c_{TW}x_3'(t) \\
M_{TW}z_3''(t) = \quad &- \quad k_{ATM}\left(\sqrt{(x_3(t)-x_1(t))^2+z(t)^2}-\sqrt{(x_3(0)-x_1(0))^2+(z(0))^2}\right)\sin(\theta) \\
&- \quad c_{ATM}\frac{d}{dt}\left(\sqrt{(x_3(t)-x_1(t))^2+z(t)^2}\right)\sin(\theta) \\
&- \quad k_{PTM}\left(\sqrt{(x_2(t)-x_3(t))^2+z(t)^2}-\sqrt{(x_2(0)-x_3(0))^2+(z(0))^2}\right)\sin(\theta) \\
&- \quad c_{PTM}\frac{d}{dt}\left(\sqrt{(x_2(t)-x_3(t))^2+z(t)^2}\right)\sin(\theta) \\
&- \quad k_{TW}z(t)-c_{TW}z'(t),
\end{aligned}
$$

with initial conditions

$$x_1(0) = 0 \text{ m}; x_2(0) = 0.00022 \text{ m}; x_3(0) = 0.00013 \text{ m}; z(0) = 0.000085 \text{ m}$$

and

$$x_1'(0) = x_2'(0) = x_3'(0) = z'(0) = 0.$$

The initial conditions were based on the precise ear measurements given in [3].

The system of equations was solved numerically using the function "ParametricNDSolve" in Wolfram Mathematica 13.3 [12].

#### S2.2 Trachea wall effective stiffness

Since the stiffness of the tree cricket trachea wall has not been empirically quantified, we used numerical parameter sweeps to determine a realistic effective stiffness ($k_{TW}$) value at 3 kHz. As a simplifying assumption, we took $k_{TW}$ to be the same in the $x-$ and $z-$ directions. For the starting point of the parameter sweeps, we used the effective stiffness of the ATM $k = 41.23$ N/m. The parameter sweeps considered the constant multiples of $k$. The constant multiples were substituted to the mathematical system outlined in Section S2.1, while keeping the remaining parameters fixed. The simulations were considered at 3 kHz, and the phase difference between the tympana was varied within the range $0° < \varphi < 180°$ for the maximum displacement magnitude of the wall.

As a benchmark for the parameter sweeps, the numerical results were compared to the Optical Coherence Tomography (OCT) recordings of the trachea wall displacement magnitude in the passive state tree cricket ear, given by Mhatre et al. [11]. The OCT recordings showed a sensitivity of $\sim$3 nm/ Pa in the $x-$direction and $\sim$4 nm/ Pa in the $z-$directions.

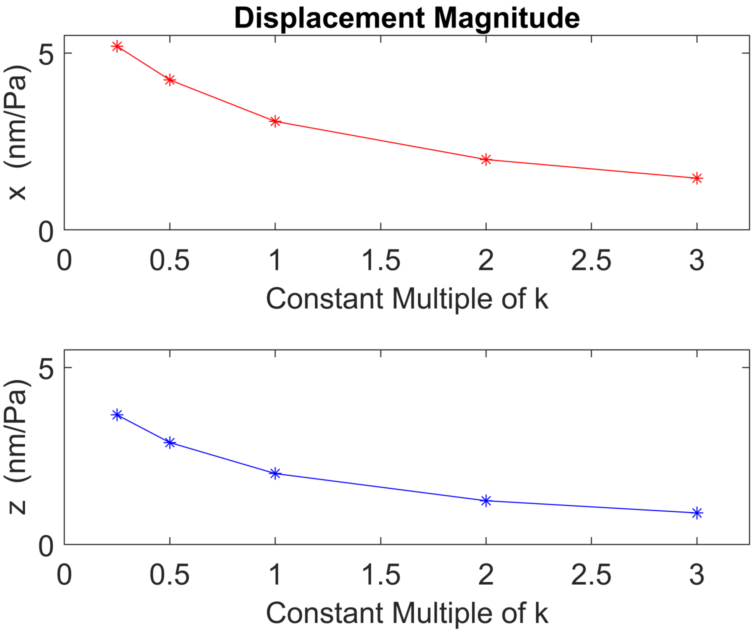


Figure S5: The numerical maximal displacement magnitude of the TW in the $x-$direction (top figure) and $z-$direction (bottom figure), for constant multiples of the effective stiffness $k$.

The numerical results for various constant multiples of $k = 41.23$ N/m are given Figure S5. Following a comparison with the experimental data, the effective stiffness of the trachea wall was adopted as $k_{TW} = \frac{1}{2}k = 20.62$ N/m.
