## Supplementary figures and images for "The tree cricket ear is a highly phase sensitive biomechanical interferometer"

### Supplemental Video

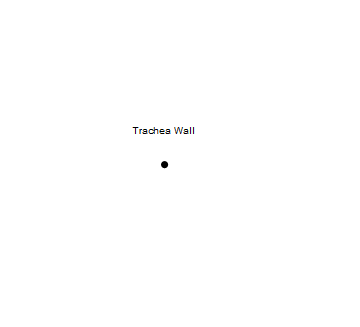
